# Adult neural stem cells and neurogenesis are resilient to intermittent fasting

**DOI:** 10.1101/2022.07.08.499318

**Authors:** Rut Gabarró-Solanas, Amarbayasgalan Davaatseren, Tatjana Kepčija, Iván Crespo-Enríquez, Noelia Urbán

## Abstract

Intermittent fasting (IF) is a promising non-pharmacological strategy to counteract ageing which has been shown to increase the number of adult-born neurons in the dentate gyrus of mice. However, it is still unclear which steps of the adult neurogenesis process are regulated by IF. The number of adult neural stem cells (NSCs) decreases with age in an activation-dependent manner. To counteract the loss of the stem cell pool, adult NSCs are mostly found in an inactive, quiescent state which ensures their long-term maintenance. We aimed to determine if and how IF impacts the activity and maintenance of adult NSCs in the hippocampus. We chose an every-other-day fasting protocol with food re-administration at night, which we found effectively induces fasting features and preserves the circadian activity pattern of mice. To determine the effects of IF on NSCs and all following steps in the neurogenic lineage, we combined fasting with lineage tracing and label retention assays. We found that IF does not affect NSC activation or maintenance. Contrary to previous reports, we also found that IF does not increase hippocampal neurogenesis. We obtained the same results regardless of strain, sex, diet length, tamoxifen administration or new-born neuron identification method. Our data suggest that NSCs maintain homeostasis upon IF and that this intervention is not a reliable strategy to increase adult neurogenesis.

## Introduction

Adult stem cells are rare cell populations in adult tissues that generate new somatic cells to support tissue turnover and repair upon injury (Li and Clevers, 2010). With age, stem cell pools decline and the function of the remaining stem cells is impaired (Goodell and Rando, 2015; López-Otín et al., 2013; Schultz and Sinclair, 2016; Tümpel and Rudolph, 2019). Fasting, either in the form of intermittent fasting or time-restricted feeding, is a dietary intervention that prolongs life and health span (Longo and Panda, 2016; Madeo et al., 2019). The health benefits of fasting are explained, in part, by improved stem cell function throughout the body, including in the intestine, muscle or hematopoietic system (Novak et al., 2021).

The adult mammalian brain harbours neural stem cells (NSCs) in specific locations such as the dentate gyrus (DG) of the hippocampus (Kuhn et al., 2018). There, NSCs are the source of adult-born granule neurons, that integrate into the pre-existing hippocampal circuit offering an additional layer of plasticity that is crucial for the modulation of memory, learning, mood or emotions (Bond et al., 2015). NSCs in the DG have a limited ability to self-renew and are mostly found in an inactive, quiescent state that prevents their activity-coupled exhaustion (Bonaguidi et al., 2011; Bottes et al., 2021; Encinas et al., 2011; Pilz et al., 2018; Urbán et al., 2016). Interestingly, NSCs can return to quiescence after proliferation, a mechanism that helps preserve the active NSC pool and sustain neurogenesis throughout life (Bottes et al., 2021; Urbán et al., 2016). Yet, the aged brain contains fewer NSCs which, in addition, shift to a deeper quiescent state resulting in a decrease in adult neurogenesis (Harris et al., 2021). Understanding NSC quiescence is therefore crucial to elucidate the mechanisms that control and support neurogenesis during adulthood and ageing.

Fasting is widely regarded as an efficient strategy to increase adult neurogenesis, therefore holding great potential to improve cognitive ability and prevent the age-related neurogenic decline (Fusco and Pani, 2013; Landry and Huang, 2021; Levenson and Rich, 2007; Mattson, 2003; Pani, 2015; Poulose et al., 2017; Van Cauwenberghe et al., 2016; Wahl et al., 2016; Zainuddin and Thuret, 2012). However, we still do not understand how NSCs and other cells along the neurogenic lineage respond to fasting. To generate new neurons, quiescent NSCs first activate and give rise to intermediate progenitor cells (IPCs). These are highly proliferative cells that amplify the progenitor pool and further differentiate into neuroblasts, eventually maturing into functional granule neurons. Fasting has been reported to promote the survival of newly born neurons (Brandhorst et al., 2015; Kim et al., 2015; Kitamura et al., 2006; Lee et al., 2002a; Lee et al., 2002b), with some works also reporting an increase in proliferation in the DG (Brandhorst et al., 2015; Cao et al., 2022; Dias et al., 2021; Park and Lee, 2011). It is therefore not clear whether fasting increases the neuronal output of adult NSCs by promoting maturation and survival of newly born neurons, by increasing proliferation of IPCs and/or by activating quiescent NSCs. A fasting-induced burst in NSC proliferation could lead to the exhaustion of the NSC pool eventually impairing adult neurogenesis. Determining the stages of the neurogenic lineage at which fasting acts to increase adult neurogenesis and whether it affects NSC activation and maintenance is necessary to predict its long-term consequences.

Here we have established an intermittent fasting (IF) protocol that effectively causes systemic changes associated with the benefits of fasting without altering the activity pattern of mice. To study the effects of IF on NSCs, we used genetic lineage tracing and thymidine analogues in combination with immunofluorescence. IF did not affect NSC proliferation or maintenance even after a 4-months long intervention. This could make IF an ideal strategy to increase adult neurogenesis without compromising the stem cell pool. However, we also found that IF did not increase adult neurogenesis at any given time. We investigated experimental variables that could explain why our results did not match previous reports on adult neurogenesis, such as mouse strain, sex, diet length, tamoxifen administration or newly born neuron quantification method. Our results consistently show that adult neurogenesis does not increase upon IF.

## Results

### Intermittent fasting with daytime refeeding disrupts the circadian activity pattern of mice

We first set out to establish a fasting protocol to study adult neurogenesis. IF is a popular form of fasting that encompasses a group of dietary interventions that rely on alternating fasting and feeding periods (Hofer et al., 2022). One of the most widely used IF interventions -and the one we chose for this study-is every-other-day-fasting (also known as alternate-day-fasting), which consists of cycles of 24h of food deprivation followed by 24h of free access to food, where food is removed or added always at the same time of the day. The specific time at which food is removed/added respect to the time at which lights are switched on in the animal house (usually referred to as Zeitgeber Time 0 or ZT0) varies among studies, although it is often not specified. Morning (ZT0-ZT4, daytime IF) and evening (ZT12-14, night-time IF) times for removing/adding food are commonly used in IF studies, including those showing effects on adult neurogenesis in young, healthy mice (M. Mattson and S. Thuret, personal communication December 2020 and June 2021 respectively). We monitored mouse behaviour in ad libitum fed (Control, free access to food), day- and night-time intermittently fasted mice for 1 month using automated home cage phenotyping PhenoMaster cages (TSE Systems). Ad libitum fed mice displayed higher activity levels during the night than during the day, with a prominent peak of activity at the beginning of the dark phase, as expected for nocturnal animals (Figure 1A, B and S1A, B). It has been previously shown that IF with daytime refeeding decouples feeding patterns from light cues and daily behaviour, flattening the rhythmic expression of circadian clock genes (Acosta-Rodríguez et al., 2022; Froy et al., 2009). Accordingly, daytime refeeding introduced an additional peak of activity during the light phase and decreased the general activity of mice during the night (Figure 1A and S1A). Night-time IF, on the other hand, synchronised food cues to the light/dark cycle preserving the normal circadian activity pattern of mice (Figure 1B and S1B). Interestingly, the total activity after 1 month of diet was not affected by day-nor night-time IF compared to control mice (Figure S1C, D). Since adult neurogenesis is influenced by circadian rhythms (Bouchard-Cannon et al., 2013; Draijer et al., 2019; Holmes et al., 2004), we chose the night-time IF protocol to study the effects of IF without introducing circadian rhythm disruption as an additional variable.

**Figure 1.**
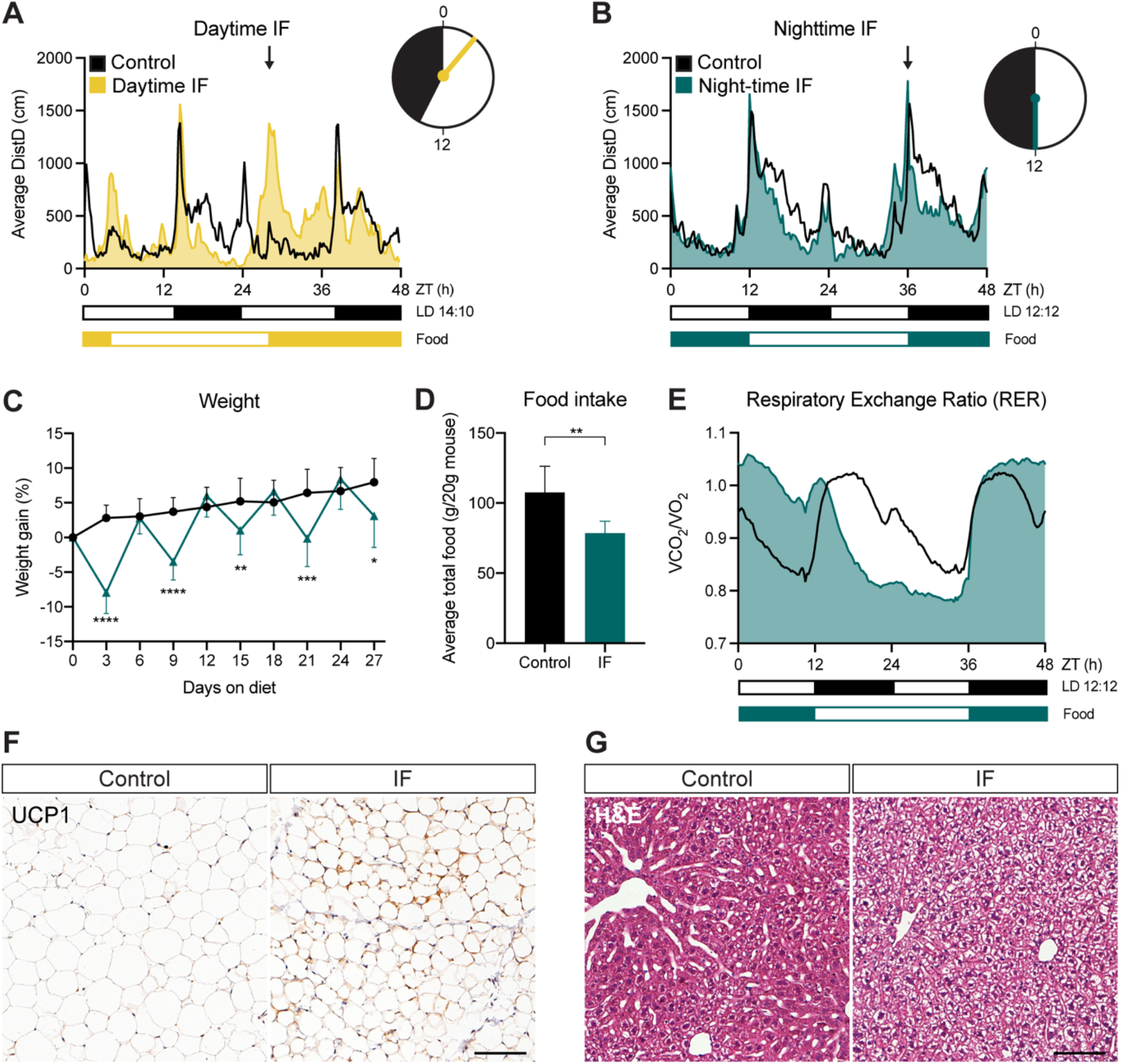
Night-time IF induces systemic features of IF without disrupting the circadian activity pattern of mice. (**A**, **B**) Mouse locomotor activity in control (ad libitum, black, n = 3 in daytime IF and n = 6 in night-time IF), daytime IF (**A**, yellow, n = 4), and night-time IF (**B**, dark blue, n = 6) mice. Activity is displayed as average distance covered by mice in 48 h cycles throughout a 1-month intervention. Black and white boxes under the graphs indicate respectively the dark and light phases of the light:dark (LD) cycle with zeitgeber time (ZT) 0 being the time of lights ON. Yellow/blue and white boxes indicate the presence and absence of food respectively. Arrows indicate the time of food availability onset. The clocks represent 24-h cycles in ZT with light and dark phases filled in white and black respectively. Clock hands indicate time of food change: ZT3 for daytime IF and ZT12 for night-time IF. Daytime IF introduces abnormal peaks of activity during the day disrupting the circadian rhythmicity of mouse activity while night-time IF preserves their normal activity pattern. (**C**) Mouse weight every 3 days shown as the percentage of weight difference to the first day of the diet. Days 3, 9, 15, 21 and 27 show weight on fasting days, and days 6, 12, 18 and 24 on feeding days. Weight oscillates in feeding and fasting days. n_control_ = 17. n_IF_ = 18. (**D**) Average total food consumed for a month shown as g of food per 20 g of mouse weight (weight reference of mice at the beginning of the diet). IF induces a mild (70%) caloric restriction. n_control_ = 17. n_IF_ = 18. (**E**) Respiratory Exchange Ratio (RER) as a comparison of CO_2_ and O_2_ volumes. RER values close to 1 indicate predominant carbohydrate metabolism as energy fuel, while RER values close to 0,7 indicate a shift towards lipid oxidation. RER is displayed as average of 48 h cycles throughout a 1-month intervention. IF induces a longer shift towards lipid oxidation. n = 6. (**F**) Representative images of UCP1-stained inguinal adipose tissue after 3 months of IF. Adipocytes of IF mice look smaller and show a mild increase in UCP1. (**G**) Representative images of liver tissue stained with H&E after 3 months of IF. IF induces hepatocyte swelling. See (**F**) and (**G**) images of all animals in **Figure S2**. Bars and error bars represent mean + SD. Significance values: *, p>0.05; **, p<0.01; ***, p<0.001; ****, p<0.0001. Scale bar: 100μm.

### Night-time IF induces characteristic features of IF in mice

We then characterised the systemic response of mice to night-time IF (from here on IF unless stated otherwise) during 1 and 3 months. Every 24h of fasting resulted in a weight loss of up to 7% of the original weight, which was recovered during the following 24h of free access to food (Figure 1C, S2A). These oscillations persisted throughout the whole 3-month intervention. We also observed that mice subjected to IF ate nearly twice as much as control mice on feeding days. Quantification of overall food consumption by the end of the treatment showed a caloric restriction of 73% or 79.8% upon 1 or 3 months of IF respectively (Figure 1D, S2B). Of note, fasted mice exhibited a similar growth curve to ad libitum fed mice (Figure 1C) and a comparable weight at the end of the IF protocol even after 3 months of diet (Figure S2A). To characterise the impact of IF on systemic metabolism we performed an indirect calorimetry during a whole month of IF to estimate the fuel source used by the body to produce energy. We used the automated home cage phenotyping system to monitor CO_2_ production and O_2_ consumption, with which we calculated the respiratory exchange ratio (RER). A RER value close to 0.7 indicates predominant usage of lipid oxidation, characteristic of resting periods, while a value around 1 indicates that carbohydrates are the main source of energy. As expected, the RER of control mice was higher during the night than during the day (Figure 1E). In IF mice, fasting extended the periods in which the body shifted its metabolism towards lipid oxidation, a feature of prolonged fasting, while feeding induced a sharp metabolic shift towards the use of carbohydrates as a fuel source (Figure 1E). Therefore, alternating fasting and feeding days resulted in the periodic metabolic shifts that are proposed to mediate the benefits of IF (Anton et al., 2018). Browning of adipose tissue is another indicator of fasting (Li et al., 2017; Liu et al., 2019). Fat browning is characterised by a reduction in adipocyte size and an upregulation of uncoupling protein 1 (UCP1), which regulates mitochondrial energy production, associated with thermogenesis in these cells (Ricquier, 2011). After 3 months of IF, inguinal adipocytes of IF mice looked smaller and had higher levels of UCP1 than those of mice fed ad libitum, although the effects were mild and variable (Figure 1F and S3). We also observed robust structural changes in the livers of IF mice (Figure 1G and S4), where hepatocytes showed an edematous morphology. This hepatocyte edema resembled the one reported upon 1-week of IF as a compensatory mechanism to fluctuating liver size between the fasting and fed states (Sarkar et al., 2021). Our data shows that liver remodelling persists after 3 months of IF. Together, these results demonstrate that cycles of 24h fasting and 24h feeding induce typical features of IF.

### IF does not affect the proliferation or maintenance of adult NSCs in the hippocampus

To specifically characterise how adult NSCs respond to IF, we used Glast-CreER^T2^;RYFP mice, a well-established mouse transgenic model for lineage tracing of NSCs in the adult DG in which Tamoxifen-induced Cre recombination indelibly labels Glast-expressing cells with YFP (Andersen et al., 2014; Mori et al., 2006; Srinivas et al., 2001). Tamoxifen was given to two-month-old mice for the 5 days prior to starting the diet and we examined the mice on a feeding day 1 month after the diet began (Figure 2A). The rate of recombination was very high throughout our experiments and identical in ad libitum-fed and intermittently fasted mice (Figure S5). We identified NSCs by their localisation in the sub-granular zone (SGZ) and the extension of a single glial fibrillary acidic protein (GFAP)^+^ radial projection to the molecular layer with help of the YFP fluorescent reporter. A small fraction of NSCs (~3.4%) was positive for the proliferation marker Ki67 in control mice, as expected for a predominantly quiescent NSC population. This number was unchanged by IF, suggesting that NSC proliferation is not affected by this intervention (Figure 2B, C). NSCs can return to quiescence after proliferating, slowing down the activation-triggered loss of the stem cell pool. We used the thymidine analogue 5-ethynyl-2’-deoxyuridine (EdU) in a label retention experiment to explore potential changes in the ability of adult NSCs to return to quiescence (Urbán et al., 2016). We administered EdU in the drinking water for 5 days to label all cells going through S-phase of the cell cycle during that period. After a 10-day chase period, we quantified NSCs that had incorporated EdU during the 5-day labelling period and had retained both the label and their NSC features (i.e. SGZ localisation and a single GFAP^+^ radial process) (Figure 2A). The percentage of EdU-retaining NSCs was similar in control and IF mice (Figure 2D, E). This, together with the similar NSC proliferation rates in control and IF mice, suggests that NSC transitions between quiescence and activation are not altered by IF. The total number of NSCs was also not changed between IF and ad libitum fed mice (Figure 2F). Together, our results show that adult NSCs are not affected by 1 month of IF.

**Figure 2.**
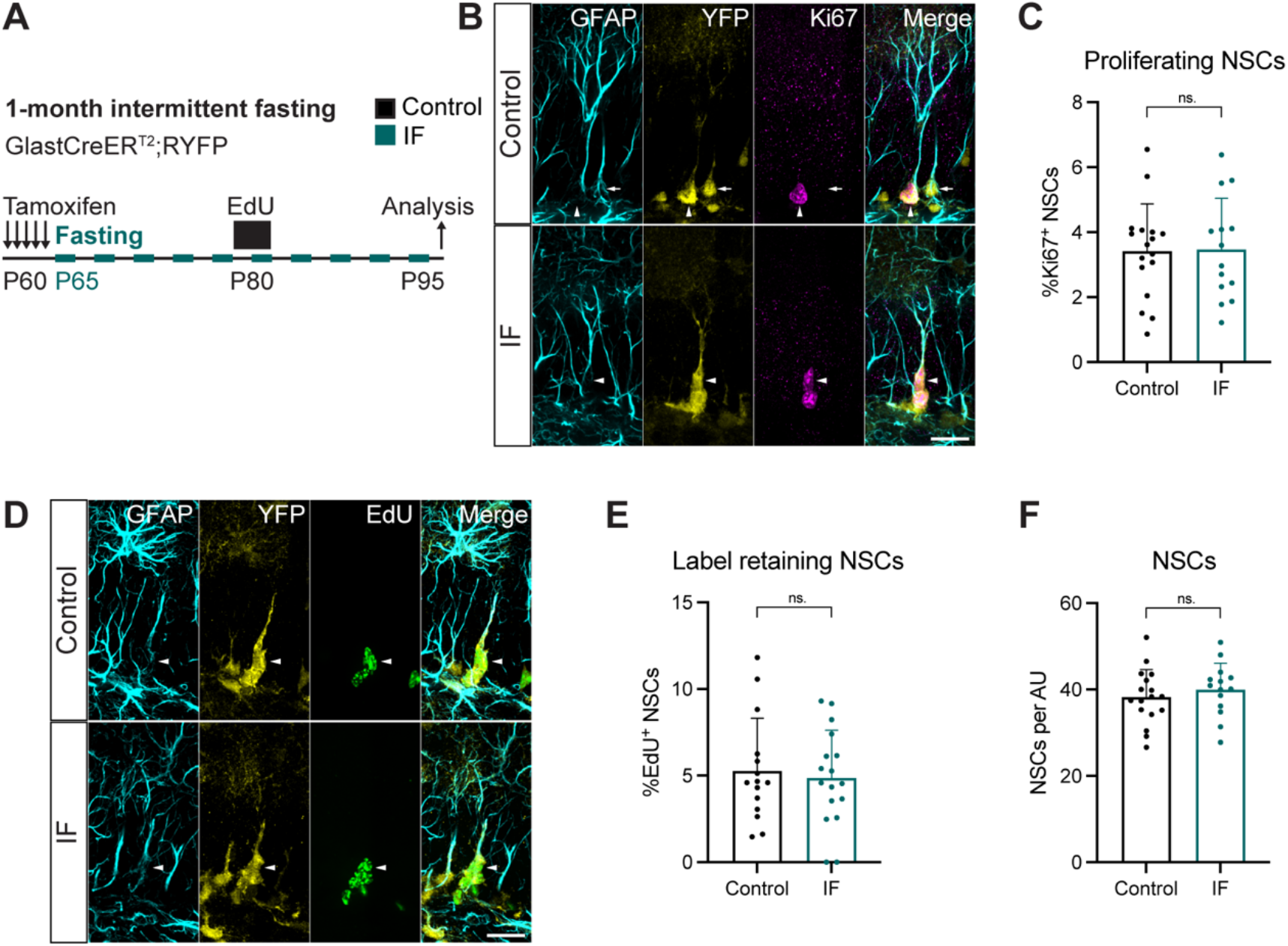
1 month of IF does not affect NSC proliferation, quiescence/activation transitions or maintenance. (**A**) 2-month-old GlastCreER^T2^;RYFP mice were administered tamoxifen on 5 consecutive days to fluorescently (YFP) label NSCs, after which, they were subjected to every-other-day IF for 1 month. EdU was administered in the drinking water for 5 days, 15 days before the analysis. (**B**, **C**) Proliferating NSCs in control and IF mice, and quantification of the percentage of proliferating NSCs. NSCs were identified by their localisation in the SGZ, the presence of a single GFAP^+^ vertical projection and the help of YFP. Nuclear colocalisation with Ki67 was used to distinguish proliferating (Ki67^+^, arrowheads) from quiescent (Ki67^-^, arrow) NSCs. The percentage of proliferating NSCs was unchanged by IF. (**D**, **E**) EdU retaining NSCs in control and IF mice and quantification of their percentage. Arrowheads indicate EdU^+^ NSCs. IF did not affect the percentage of label retaining NSCs. (**F**) Total number of NSCs normalised to DG length. The number of NSCs is unaffected by IF. Bars and error bars represent mean + SD; dots represent individual mice. Significance values: ns, p>0.05. Scale bar: 20μm. AU: arbitrary unit. Black: control (ad libitum), dark blue: IF.

To understand the consequences of longer exposures to IF, we subjected Glast-CreER^T2^;RYFP mice to 3 months of IF (Figure 3A), which has been shown to be enough to increase neuronal production in the DG (Baik et al., 2020; Dias et al., 2021; Kim et al., 2015; Kitamura et al., 2006; Lee et al., 2002b; Lee et al., 2002a; Li et al., 2020). The percentage of proliferating NSCs in the 3-month intervention was halved compared to the same value in the 1-month intervention, in accordance with the reported increase in quiescence during ageing (Harris et al., 2021) (Figures 2B, C and 3B). NSC proliferation did not change upon IF (Figure 3B). We then examined NSC maintenance by quantifying the total number of NSCs present at the end of the treatment (Figure 3C, D). Despite the well-established decrease of neurogenesis with age, we did not observe a decrease in total NSCs in 5-month-old mice respect to 3-month-old mice (Figures 2F and 3C). This could be explained by batch variability in the stainings or by a shift in the exponential phase of the decline in NSC numbers to either before 3 months or after 5 months of age in these mice. Nevertheless, the number of NSCs in IF mice were indistinguishable from those of ad libitum fed mice (Figure 3C, D).

**Figure 3.**
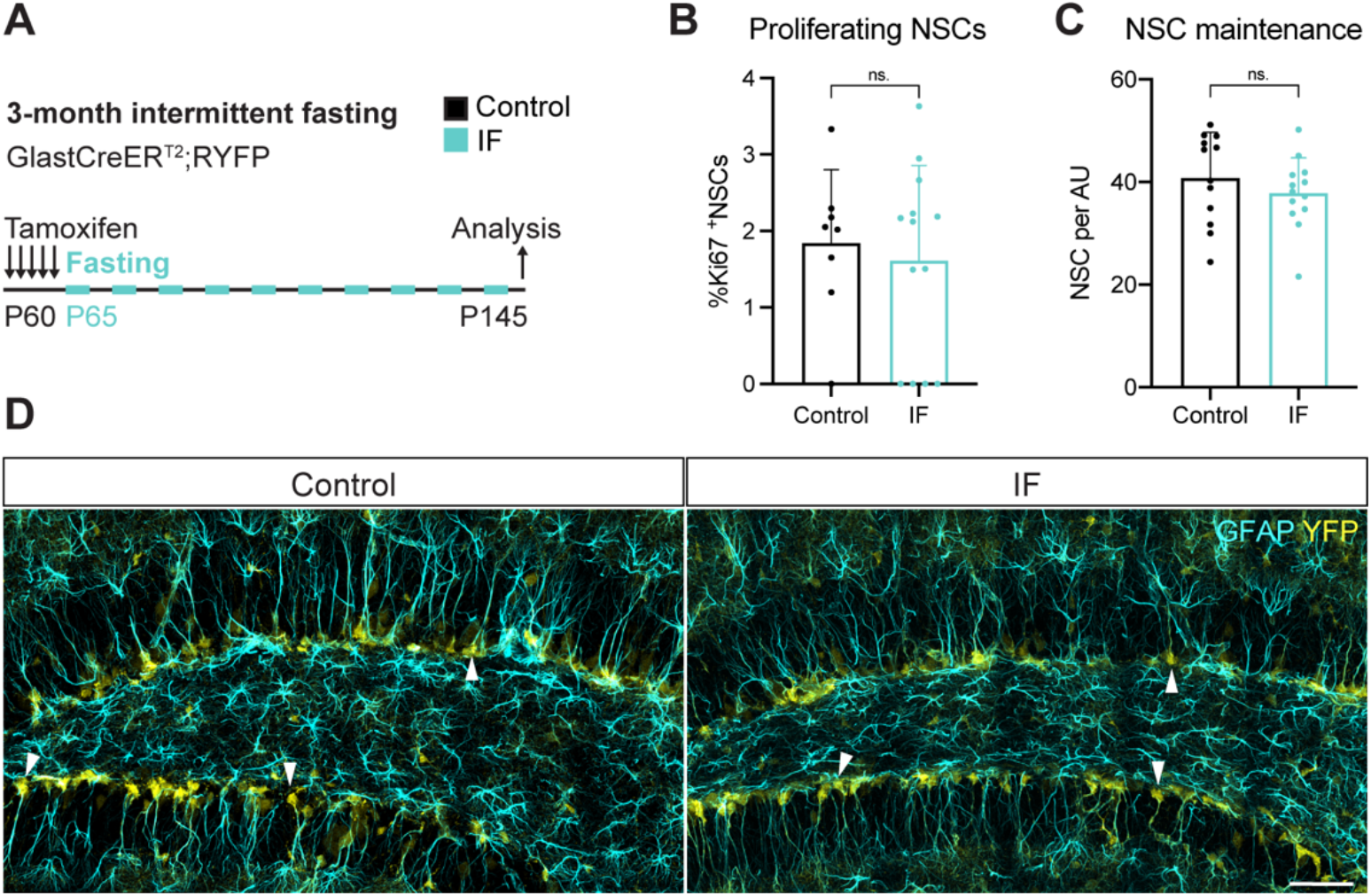
3 months of IF do not affect NSC proliferation nor maintenance. (**A**) 2-month-old GlastCreER^T2^;RYFP mice were administered tamoxifen on 5 consecutive days to fluorescently (YFP) label NSCs, after which, they were subjected to every-other-day IF for 3 months. (**B**, **C**) Quantification of the percentage of proliferating NSCs and the total number of NSCs in control and IF mice normalised to DG length. (**D**) NSCs identified by their localisation in the SGZ, the presence of a single GFAP^+^ vertical projection, and the help of YFP. Arrowheads show several NSCs. IF mice show similar levels of proliferating and total NSCs to control mice. Bars and error bars represent mean + SD; dots represent individual mice. Significance values: ns, p>0.05. Scale bar: 50μm. AU: arbitrary unit. Black: control (ad libitum), light blue: IF.

These data indicate that NSCs are protected from proliferation bursts that could lead to exhaustion, making IF a safe option for potentially increasing neurogenesis through diet-based interventions.

### Adult neurogenesis is mildly and transiently impaired by IF

Since neither the pool nor the activity of NSCs change upon IF, we hypothesized that the previously reported increase in neurogenesis laid in subsequent stages of neurogenesis. We set out to determine at which step/s of the neurogenic lineage IF influences adult neurogenesis. NSCs and neuroblasts are predominantly out of the cell cycle, meaning that the vast majority of proliferating cells in the SGZ are IPCs (Hodge et al., 2008; Kronenberg et al., 2003). We therefore used Ki67 as a proxy for IPC identification. As previously reported, we observed a decrease in IPC abundance over time (Figure 4A, B). IF neither changed the number of IPCs after 1 month nor prevented their decline after 3 months (Figure 4A, B).

**Figure 4.**
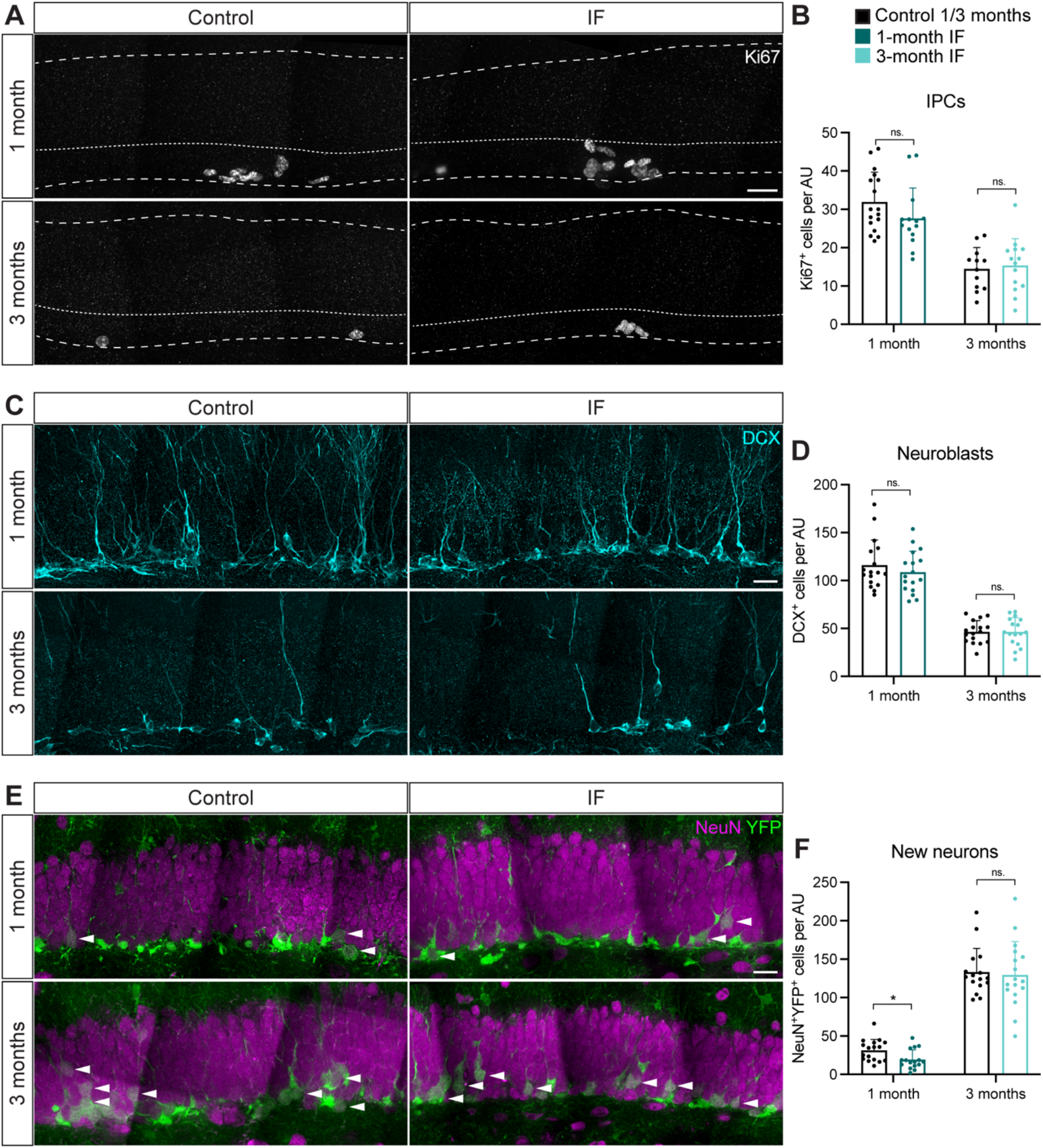
IF does not affect IPCs nor neuroblasts and only mildly and temporarily decreases the neuronal output. (**A**, **B**) Ki67^+^ cells in control and IF mice after a 1-or 3-month intervention as a proxy for IPCs and quantification of IPCs normalised to DG length. Dashed and dotted lines indicate DG area and border between the SGZ and the molecular layer respectively. IPCs were unchanged by IF. (**C**, **D**) Neuroblasts identified by the marker for immature neurons DCX and quantification. IF did not affect the number of neuroblasts. (**E**, **F**) Newly born neurons (arrowheads) identified by the colocalization of the mature neuronal marker NeuN and the YFP reporter, indicating that these neurons were generated from the population of NSCs labelled with YFP at the beginning of the diet. Quantification of newly born neurons normalised to DG length. The number of new neurons transiently decreased upon 1 month of IF, but was unchanged after 3 months. Bars and error bars represent mean + SD; dots represent individual mice. Significance values: ns, p>0.05; *, p<0.05. Scale bar: 20μm. AU: arbitrary unit. Black: control (ad libitum) for each experiment, dark blue: 1-month IF, light blyue: 3-month IF.

Next, we quantified the number of neuroblasts, identified by doublecortin (DCX) immunoreactivity. Our results show that neuroblasts were more abundant in younger mice and that they are generated at the same rate in IF mice and ad libitum fed mice at both time points (Figure 4C, D).

Finally, we measured the number of new neurons generated during the intervention. We used colocalization of the YFP reporter and the marker for mature neurons NeuN to identify newly born neurons. In accordance with the cumulative labelling of neurons upon lineage tracing, mice had many more newly born neurons labelled with the YFP reporter after 3 months than after 1 month (Figure 4E, F). Unexpectedly, we found a reduction in the number of newly born neurons in IF mice compared to ad libitum fed mice after 1 month of IF (Figure 4E, F). Cell death, measured by the number of picnotic nuclei in the SGZ, was comparable in control and IF mice (Figure S6A). Since neuroblasts were produced at the same rate (Figure 4D), we reasoned that this decrease in neurogenesis could be due to a transient drop in neurogenesis at an earlier time point. We made use of the label retention experiment in Figure 2A to ask if neuronal production was impaired two weeks after starting IF. Indeed, we found that fewer neurons had been produced at that time in mice subjected to IF compared to control mice (Figure S6B). One month of IF is therefore not only insufficient to increase adult neurogenesis but in fact decreases the neuronal output of adult NSCs. The decrease of neurogenesis was transient, as the number of newly born neurons went back to control levels in the 3-month intervention (Figure 4E, F), suggesting that the process of adult neurogenesis is robust enough to buffer initial changes in neuronal production.

Our findings show that 1 or 3 months of IF do not increase adult neurogenesis. This seemingly contradicts previous reports and puts a question mark on the suitability of IF as a tool to increase neurogenesis in the adult hippocampus.

### IF does not increase adult neurogenesis regardeless of sex, labelling method, strain, tamoxifen usage or diet length

We next asked which intrinsic variables of our study could explain the contradicting results with previously published studies.

The metabolism of male and female mice has been shown to react in a sex-specific manner to fasting and refeeding (Freire et al., 2020). While most studies on IF and neurogenesis were conducted with only male or only female mice, differences in the response to caloric restriction of adult hippocampal neurogenesis have been reported between sexes (Park et al., 2013). In our experimental set up, we used enough males and females to identify sex differences in the response of adult neurogenesis to IF. We found that the transient decrease in newly born neurons after 1 month of IF was affected by sex (Figure 5A). However, we did not observe sex differences at any other point of the study, with neurogenesis not being affected by IF in either male or female mice (Figure 5B and S7). Of note, data variability was also similar in male and female mice (Figure S7).

**Figure 5.**
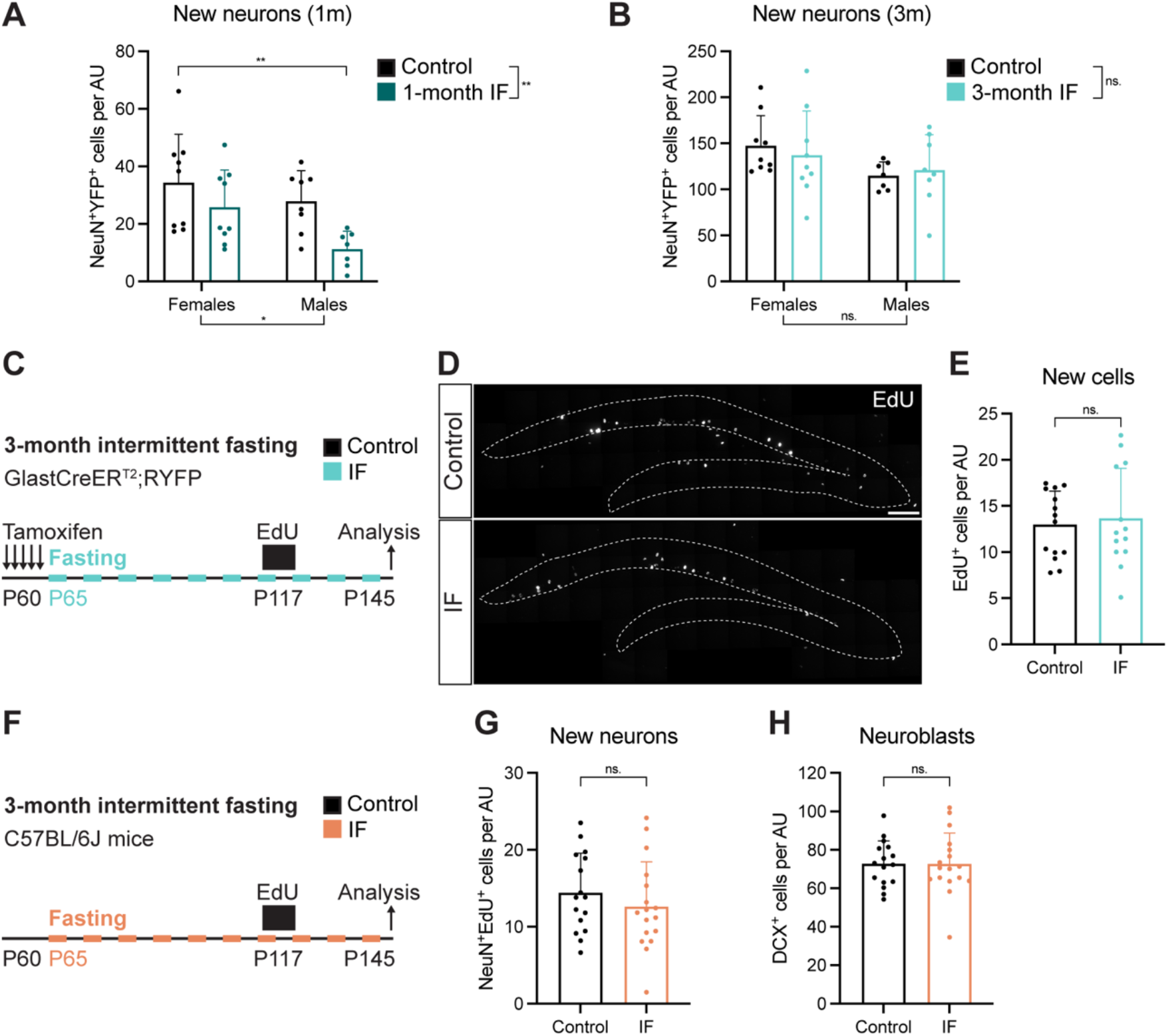
Sex, labelling method, tamoxifen or strain do not alter the neurogenic response to IF. (**A**, **B**) Quantification of newly born neurons after 1 (**A**) or 3 (**B**) months of IF segregated by sex. Significance signs outside the graph indicate result of a two-way ANOVA and sign in the graph indicates result of a Tuckey’s multiple comparison test. See all sex-segregated data in **Figure S5**. Sex differences were found only in the number of new neurons at 1 month. (**C**) Survival assay with a thymidine analogue to replicate the labelling method of previous publications. EdU was administered for 4 days to GlastCreER^T2^;RYFP mice (see experiment in **Figure 3**) that had undergone 2 months of IF to label a cohort of proliferating cells that was analysed after a 1 month chase, giving the cells time to progress in the neurogenic lineage. (**D**, **E**) New cells in the DG (enclosed in dashed lines) shown by EdU-labelled cells 1 month after EdU administration. The number of new cells in the DG was unchanged by IF. (**F**) Evaluation of tamoxifen and strain impact in the effects of IF on adult neurogenesis. 2-month-old C57BL/6J mice were subjected to 3 months of IF. A survival assay with a thymidine analogue 1 month before the end of the diet was included. (**G**) Quantification of newly born neurons identified by colocalization of EdU and the marker for mature neurons NeuN. (**H**) Quantification of new neuroblasts. IF did not increase the number of new neurons or neuroblasts in C57BL/6J mice. Bars and error bars represent mean + SD; dots represent individual mice. Significance values: ns, p>0.05; *, p<0.05; **, p<0.01. Scale bar: 100μm. AU: arbitrary unit. Black: control (ad libitum) for each experiment, light blue: 3-month IF, orange: 3-month IF C57BL/6J.

In this study, we used genetic lineage tracing to quantify the total number of new neurons generated from NSCs throughout the whole dietary intervention. Instead, previous studies relied on the use of thymidine analogues (BrdU) to label a cohort of proliferating cells and quantify the amount of total BrdU^+^ cells or BrdU^+^ neurons 1 month after labelling, obtaining information exclusively on one neurogenic wave at the end of the treatment (Brandhorst et al., 2015; Dias et al., 2021; Kim et al., 2015; Kitamura et al., 2006; Lee et al., 2002b; Lee et al., 2002a). To assess the effects of IF in a comparable way, we used EdU to pulse-label proliferating cells 1 month before the end of the 3-month IF protocol (Figure 5C). Again, the number of EdU^+^ cells (Figure 5D, E) was unchanged by the diet, showing that IF does not affect neurogenesis after three months.

Another particularity of our study is that the GlastCreER^T2^;RYFP mice we used had a mixed genetic background (Austin et al., 2020), while previous studies had used mice from the C57BL/6J strain. Strain-specific differences in the response of adult neurogenesis to dietary interventions such as high-fat diet have been reported (Hwang et al., 2008). In addition, recurrent doses of Tamoxifen alter neurogenesis and induce broad detrimental effects, from damaging the gut epithelia to impairing microglial reactivity (Smith et al., 2022). To test whether strain and tamoxifen could alter how IF modulates adult neurogenesis, we subjected C57BL/6J mice to 3 months of IF, excluding tamoxifen administration (Figure 5F). We did a 1-month EdU-chase, as in our previous experiment with Glast-CreER^T2^;RYFP mice, and analysed the number of EdU^+^ cells at the end of the treatment. We identified newly born neurons as EdU^+^NeuN^+^ cells (Figure 5G) and, again, we did not find any differences caused by the diet. In addition to that, neuroblasts (Figure 5H) were not altered by IF either suggesting that tamoxifen and strain do not explain the discrepancy in results.

Lee et al., Kim et al., and Dias et al. reported an increase in neurogenesis in a cohort of cells that was labelled at 3 months of IF and analysed 1 month later, 4 months after the start of the diet (Dias et al., 2021; Kim et al., 2015; Lee et al., 2002a). Since we were interested in determining whether NSCs were at the origin of this increase, we performed 3-month-long experiments, meaning that neuronal production was also analysed after 3 months of IF. Therefore, we next asked whether extending the diet for an additional month would increase neurogenesis. We used male C57BL/6J mice and labelled a cohort of proliferating cells with EdU for 12 days, 3 months after the start of the diet (Figure 6A). We analysed newly born neurons 1 month after, and, to our surprise, observed a decrease in EdU^+^NeuN^+^ cells (Figure 6B, C). We wondered if this was a result of decreased NSC activation, as a mechanism to prevent NSC exhaustion and prolong neurogenesis during aging, but IF did not change NSC proliferation nor maintenance (Figure 6D, E). Other stages of neurogenesis were also unaffected since there were the same number of IPCs and neuroblasts in control an IF mice (Figure 6F, G).

**Figure 6.**
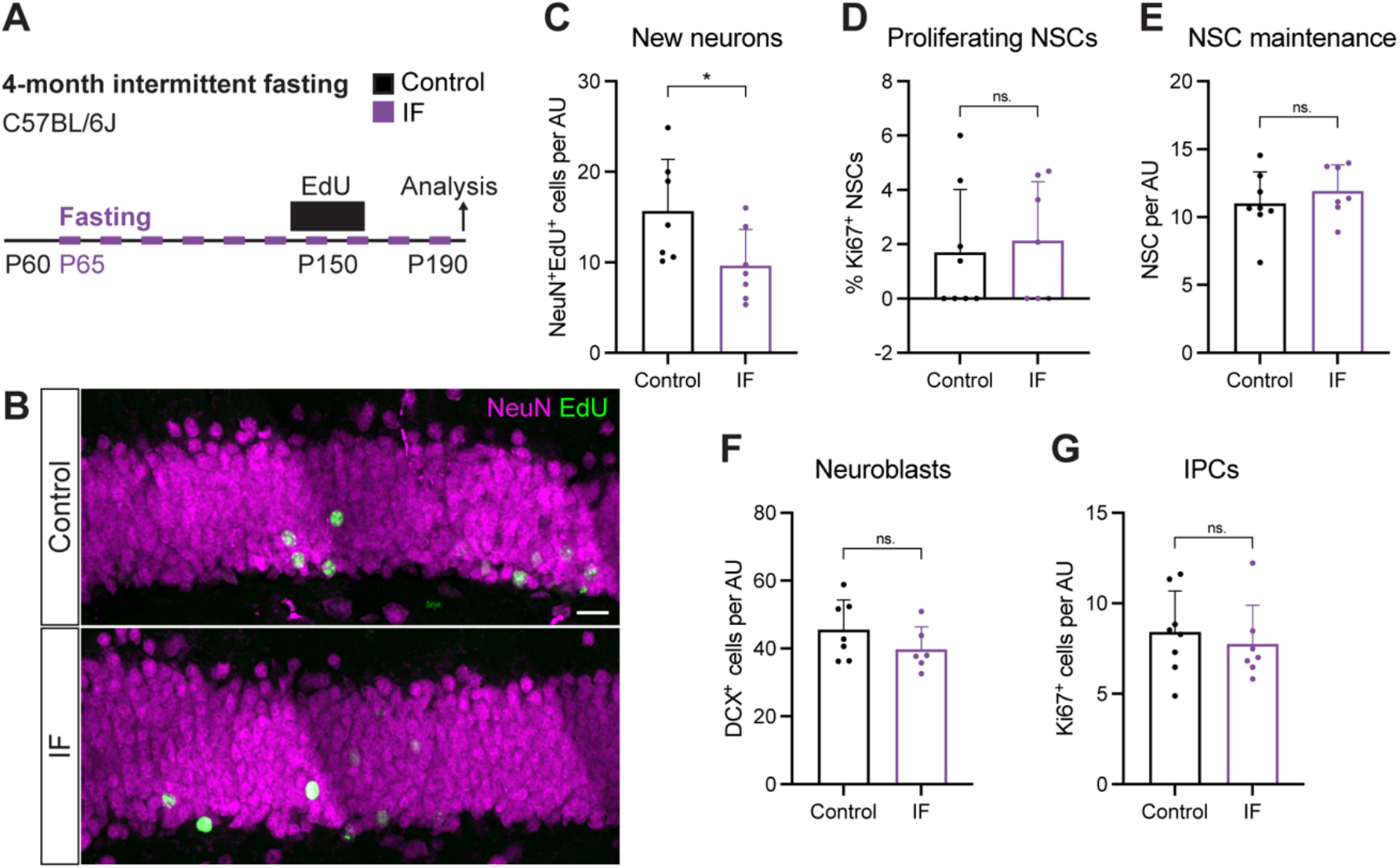
4 months of every-other-day IF do not increase adult neurogenesis. (**A**) 2-month-old male C57BL/6J mice were subjected to 4 months of IF, as in previous studies that reported increases in adult neurogenesis. A survival assay with a thymidine analogue (12-day labelling - 1 month chase) was used to evaluate neurogenesis. (**B**, **C**) Image and quantification of newly born neurons identified by colocalization of EdU and the marker for mature neurons NeuN. 4 months of IF induce a decrease in EdU-labelled neurons. (**D**) Quantification of proliferating NSCs. (**E**, **F**, **G**) Quantification the total number of NSCs, neuroblasts and IPCs normalised to DG length. The neurogenic lineage is not affected by 4 months of IF. Bars and error bars represent mean + SD; dots represent individual mice. Significance values: ns, p>0.05; *, p<0.05. Scale bar: 20μm. AU: arbitrary unit. Black: control (ad libitum), purple: IF 4-months C57BL/6J.

Together, these results show that differences in sex, labelling method, strain, tamoxifen usage and diet length do not explain the discrepancies in the response of adult neurogenesis upon IF between our study and others. We conclude that every-other day IF is not a robust strategy to increase adult neurogenesis in the hippocampus.

## Discussion

Fasting is a promising non-pharmacological intervention to increase life span and counteract ageing that has been associated with increased adult neurogenesis. However, few studies have investigated the specific effects of fasting on adult NSCs and other adult stem cell populations. This is important not only because impaired adult stem cell function is regarded as one of the hallmarks of ageing, but also because alterations in the adult stem cell population will have long-lasting consequences for tissue homeostasis. Here, we used genetic lineage tracing to determine the effects of every-other-day IF on adult NSCs.

We found that IF does not affect the proliferation or maintenance of adult NSCs. This suggested that NSCs are preserved upon IF and prompted us to investigate which further steps of neurogenesis might be increased by fasting. We carefully monitored the progression of the neurogenic lineage upon 1 or 3 months of IF and found no differences between control and fasted mice. This is in contrast with previous work reporting a sharp increase in proliferative cells (BrdU labelled) in the DG after 3 months of every-other-day IF (Dias). One possible explanation is that Dias et al. analysed the mice after 5 days of a recovery period with ad libitum feeding, while our mice were still fasting at the time of analysis. Alternating fasting periods with normal feeding has been proposed to be a crucial mechanism by which IF and CR exert their beneficial effects (Brandhorst et al., 2015). It would be very interesting to determine whether returning to ad libitum feeding from IF rather than fasting on its own can promote cell proliferation along the neurogenic lineage. However, a recent report using a similar intervention consisting of two fasting days per week (known as 5:2 diet) shows no effect on proliferation in the DG, similar to our results, even after a refeeding period (Roberts et al., 2022).

Our results were therefore in line with numerous reports which did not find differences in cell proliferation upon fasting (Kitamura et al., 2006; Lee et al., 2002b; Lee et al., 2002a). Those studies found instead an increased survival of newly born cells 4 weeks after fasting. We used lineage tracing to quantify the neuronal output of hippocampal NSCs and found no difference between control mice and mice fasted for 3 months. Classically, the effects of IF on adult neurogenesis were evaluated through neuronal survival assays with thymidine analogues (Kim et al., 2015; Lee et al., 2002a). Thymidine analogues (such as BrdU, CldU, IdU or EdU) are administered to the mice to label proliferating cells, with neurogenesis being evaluated after a chase period by direct quantification of the amount of newly born cells and/or comparison to the initially labelled population. This method evaluates only narrow waves of neurogenesis and, when several time points are used, can be useful to identify changes in the effects of the intervention over time. However, it also increases the variability of the results and, unlike lineage tracing, does not provide information on the total number of new neurons produced during the whole intervention. We nevertheless performed a classical label retention experiment to our mice and again found no increase in neuronal production upon IF for 3 or 4 months. Of note, we observed high variability in the effects of IF on adult neurogenesis, with an initial decrease followed by no effect and a final further decrease in newly born cells. The effects we observed were small and only emerged when analysing small cohorts of new neurons, either using label retention or short (1-month) lineage tracing. They could therefore be a consequence of the high degree of intrinsic variability in adult neurogenesis, questioning their functional importance. Since other cells proliferate in the dentate gyrus and can uptake the label (e.g. astrocytes, oligodendrocyte precursors or microglia), the interpretation of label-retention data is challenging unless combined with neuronal markers. In addition, the thymidine analogue positive population by the end of the labelling period depends not only on uptake of the label by proliferating cells but also on their differentiation and survival (Kronenberg 2003), hampering the interpretation of the results. By using lineage tracing combined with immunofluorescence and thymidine analogues in label-retention experiments, we could separately evaluate different steps along the neurogenic lineage and faithfully quantify changes - or the lack of them - in neurogenic output.

To perform lineage tracing of adult NSCs, we used GlastCre mice that have a mixed genetic background. However, C57BL6/J is the most commonly used strain in fasting studies (REFS). In accordance with different strains presenting different levels of adult neurogenesis (Goodrick et al., 1990; Kempermann et al., 2006; Kim et al., 2017; Koehl, 2015; Wiget et al., 2017), we did observe differences in the number of neuroblasts between control GlastCre mice and control C57BL/6J at 5 months of age (Figure S8). Therefore, we repeated our experiments using C57BL/6J mice. Our results again showed that 3 or 4 months of fasting did not increase neurogenesis in the hippocampal niche. However, since variations can occur even within the C57BL/6J strain and the response to dietary interventions is strain and even substrain-specific (Bachmann et al., 2022; Hwang et al., 2008; Mitchell et al., 2016; Siersbæk et al., 2020), we cannot rule out that small differences in the animals’ background could have contributed to the discrepancies between previous studies and ours.

Differences in the metabolic response to dietary interventions have also been observed between male and female mice (Bachmann et al., 2022; Mitchell et al., 2016), making it crucial to include both sexes when analysing the effects of any potential clinically relevant interventions. Although increased neurogenesis upon fasting has been reported for both sexes, we wondered whether differences in basal levels and response to the diet could be masking the effects of fasting in our data. Splitting our data by sex showed no increase in adult neurogenesis upon fasting in either male of female mice. We observed one parameter that was affected by sex: the number of new neurons after 1 month of IF (Figure 5A, 5B and S7). Since sex differences are particularly important for stress-related factors (Yagi and Galea, 2019), we hypothesize this discrepancy could be linked to metabolic or stress-related differences in handling the change from ad libitum feeding to fasting between male and female mice. Of note, while female mice are often excluded arguing higher variability in the results, variability was comparable in both sexes in our data (Figure S7).

Although some reports indicate that weight loss is not necessarily linked to the benefits of IF and CR, it remains plausible that the degree of caloric restriction and weight loss induced by IF could influence its subsequent effects on adult neurogenesis. Both Lee et al. and Kim et al. reported weight loss upon every-other-day IF. Our protocol, in contrast, did not affect the final weight of the mice or their growth curve. This is despite a reduction in calorie consumption of more than 20% and the presence of clear weight oscillations between fasting and feeding days. Our mice also showed signs of broad systemic changes induced by fasting (inguinal adipose tissue browning and liver remodelling), but we cannot rule out that these were not strong enough to induce changes in adult neurogenesis.

While establishing our every-other-day IF protocol, we found that daytime IF disrupts the circadian activity pattern of mice, introducing a confounding variable to IF that could affect adult neurogenesis (Bouchard-Cannon et al., 2013; Draijer et al., 2019; Schouten et al., 2020). A very recent report supports this notion, as circadian entraining can boost the beneficial effects of caloric restriction on longevity (Acosta-Rodríguez et al., 2022). This calls for a close monitoring of potential circadian disruptions during fasting interventions and for careful reporting of fasting protocols. However, circadian disruption alone does not explain the differences between our results and previous studies, as both day and night-time IF have been shown to increase neurogenesis in young, healthy mice (M. Mattson and S. Thuret, personal communication).

There are multiple other variables that we did not test for in our study which can modulate the impact of fasting on neurogenesis. These include mice stress and overall fitness (Snyder et al., 2011; Toda et al., 2019), pathological states (Cao et al., 2022; Li et al., 2020; Wu et al., 2008), age (Brandhorst et al., 2015; Park et al., 2013) and even cage bedding (Gregor et al., 2020) and diet composition (Cryan and Dinan, 2012; Ribeiro et al., 2020; Wei et al., 2021). We also did not assess differences in behavioural performance in our mice, which could be very interesting to determine the involvement of adult neurogenesis in the cognitive benefits of fasting. Of note, a recent report showing no effect on adult neurogenesis upon a 5:2 fasting intervention in mice also showed a lack of cognitive improvement in those mice (Roberts et al., 2022).

In summary, the specific combination of sex, strain, health status, housing conditions and fasting protocol can influence the outcome of dietary interventions on adult neurogenesis, putting a question mark on IF as a strategy to boost neurogenesis. Despite this, diet is regarded as an important regulator of adult neurogenesis (Aimone et al., 2014; Denoth-Lippuner and Jessberger, 2021; Urbán et al., 2019), with fasting often highlighted as a positive modulator of neuronal production (Bouchard and Villeda, 2015; de Lucia et al., 2017; Fusco and Pani, 2013; Katsimpardi and Lledo, 2018; Landry and Huang, 2021; Levenson and Rich, 2007; Mattson, 2003; Murphy et al., 2014; Pani, 2015; Park and Lee, 2011; Poulose et al., 2017; Valero et al., 2016; Van Cauwenberghe et al., 2016; Wahl et al., 2016; Zainuddin and Thuret, 2012). The effects of fasting on adult neurogenesis are, however, very diverse, ranging from variable positive impact on the number of newly generated neurons (Apple et al., 2019; Baik et al., 2020; Brandhorst et al., 2015; Cao et al., 2022; Dias et al., 2021; Kaptan et al., 2015; Kim et al., 2015; Kitamura et al., 2006; Kumar et al., 2009; Lee et al., 2000; Lee et al., 2002b; Lee et al., 2002a; Li et al., 2020; Park et al., 2013; Wu et al., 2008) to no effect or even a mild impairment (Bondolfi et al., 2004; Roberts et al., 2022; Staples et al., 2017). We propose the variability is due to a combination of different factors (and most likely different combinations of factors in each case). This fits with a model in which IF can render NSCs and other cells along the neurogenic lineage more susceptible to further interventions such as circadian disruption or sudden re-feeding (Urbán, 2022). In addition, publication bias might be playing a role in skewing the literature on fasting and neurogenesis towards reporting positive results. In some reviews, even studies reporting no effect are cited as evidence for improved neurogenesis upon IF. Reporting of negative results, especially those challenging accepted dogmas, and a careful and rigorous evaluation of the publications cited in reviews are crucial to avoid unnecessary waste of resources and to promote the advancement of science.

## Materials and Methods

### Animal husbandry

Mice were housed in groups in ventilated cages (TECNIPLAST 1285L NEXT) provided with HEPA filtered air covered with Aspen-wood-derived bedding (Kliba Nafag, ABEDD 4063 OM G10). They were kept under a 12:12 h light/dark cycle (light period 0600 – 1800 h CET), unless stated otherwise. Mice had ad libitum access to food (Ssniff, V1184-300) and water. To generate a mouse line that allowed lineage tracing of adult NSCs with a fluorescent reporter (Andersen et al., 2014), we crossed Glast-CreER^T2^ (Slc1a3^tm1(cre/ERT2)Mgoe^, (Mori et al., 2006)) mice and RYFP (Gt(ROSA)26Sor^tm1(EYFP)Cos^, (Srinivas et al., 2001)) mice. All Glast-CreER^T2^;RYFP mice were of mixed genetic background. 4–6-week-old C57BL/6J (Charles River) mice were obtained from the Comparative Medicine facility at IMBA/IMP. Experimental groups were formed by randomly assigning mice from different litters within each mouse strain and all experiments were conducted in male and female mice, except stated otherwise.

All animals were bred and maintained in accordance with ethical animal license protocols complying with Austrian and European legislation.

### Intermittent fasting

Mice were randomly divided into Control or IF groups at postnatal day 65 (P65). Control mice had ad libitum access to food. Mice in the IF group were subjected to an every-other-day fasting regime, comprising 1 day of fasting followed by 1 day of feeding. For night-time IF, food was removed at 1800 h CET coinciding with the time at which the lights went off (ZT12) and reintroduced at 1800 h CET on the following day (after 24 h). For daytime IF, food change happened every day at 0900 h CET (ZT3). Mouse weight was recorded every 3 days in both Control and IF groups and leftover food weighted once per week. Weight gain was calculated as the percentage of the weight difference to the weight at the beginning of the diet (P65). Total food intake was calculated by the difference between supplied and leftover food over the treatment in each cage and divided by the number of animals per cage. The length of each experiment is specified in the text and figure legends. For the 4-months-long experiment depicted in Figure 6, mice were fed automatically on a defined schedule through an internally developed automated system.

### Automated home cage mouse phenotyping

Locomotor and metabolic activity were measured using the PhenoMaster system TSE Systems every 15 min during a month. Mice were acclimated to PhenoMaster water bottles for at least 6 days and to the complete PhenoMaster housing (TECNIPLAST GreenLine GM 500) for 2 days. Mice in the daytime IF experiment were housed in groups (2-3 mice per cage) in a 14:10 h light/dark cycle (light period 0600-2000 h CET). Mice in the night-time IF experiment were housed individually in a 12:12 h light/dark cycle (light period 0600-1800 h CET). Cages were kept in a climate chamber in the same environmental conditions as the general animal rooms (24°C and 50% humidity). The PhenoMaster cages were programmed to automatically grant or restrict access to the food basket at the designated time.

Before each run, CO_2_ and O_2_ sensors were calibrated against a defined mix of CO_2_ and O_2_. The respiratory exchange ratio (RER) was calculated as the ratio of VCO_2_ produced to the VO_2_ consumed by the PhenoMaster software. Mouse locomotor activity was monitored using the ActiMot2 Activity module equipped with 3-dimensional infrared sensors. An estimate of movement (DistD, cm) was calculated using the PhenoMaster software. Data acquired in 15-min intervals are expressed as means of 48h cycles. For grouped housed mice (daytime IF), DistD was divided by the number of mice in each cage. Total mouse locomotor activity was calculated as the sum of cm throughout the whole intervention and segregated by time periods to depict differences between diurnal and nocturnal behaviour during fasting and feeding days. For daytime IF, where the mice were kept in a 14:10 LD cycle, the data was divided in ZT 0-14-24-38-48 h. For night-time IF, where the mice were kept in a 12:12 LD cycle, the data was divided in ZT 0-12-24-36-48 h.

The automated home cage mouse phenotyping was performed by the Preclinical Phenotyping Facility at Vienna BioCenter Core Facilities (VBCF), member of the Vienna BioCenter (VBC), Austria.

### Tamoxifen and EdU administration

To induce activation of CreER^T2^ recombinase, Glast-CreER^T2^;RYFP mice were intraperitoneally injected with 2 mg (75-100 mg/kg) of Tamoxifen (Sigma, T5648) diluted in corn oil (Sigma, C8267) at postnatal day 60 (P60) for 5 consecutive days. Recombination levels were evaluated by comparing YFP^+^ neuroblasts (DCX^+^ cells) to the total number of neuroblasts (Figure S5). Recombination was very high in both experiments (1 month and 3 months - above 90%) and levels were comparable in Control and IF groups.

To label cells in S-phase, 5-ethynyl-2’-deoxyuridine (EdU, Carl Roth, 7845) was administered in drinking water (0,1 mg/mL) ad libitum. The length of EdU administration and chase period is specified in the text and figures.

### Tissue preparation and immunohistochemistry

Mice were transcardially perfused with phosphate buffered saline (PBS) for 3 min, followed by 4% paraformaldehyde (PFA, Sigma, P6148) in PBS for 15 min.

Brains were post-fixed in 4% PFA for 2-6 h at 4°C with rocking, washed with PBS and stored in PBS with 0.02% Sodium Azide (ITW Reagents, A1430). Brains were coronally sectioned at a thickness of 40 μm using a vibratome (Leica, VT1000S). Immunofluorescence of brain tissue was performed in free-floating sections. Sections were blocked in 1%Triton-PBS with 10% normal donkey serum (DS, Jackson Immuno Research, JAC0170001210) for 2 h at room temperature. Primary and secondary antibody solutions were prepared in 0.1%Triton-PBS with 10% DS. Sections were incubated in primary antibody solution at 4°C overnight, washed 3 × 15 min with 0.1%Triton-PBS and incubated with secondary antibody solution at room temperature for 2 h. Following 1 × 0.1%Triton-PBS and 2 × PBS 15 min washes, sections were incubated with 1 μg/mL DAPI (Sigma, D9542) in 1:1 PBS:H2O at room temperature for 30 min. EdU was detected before DAPI incubation using Click-iT™ Plus Alexa Fluor™ Picolyl Azide tool kits with 488 and 647 dyes (Invitrogen, C10641 and C10643 respectively). All incubations and washes were performed under gentle rocking.

Inguinal adipose tissue and livers were dissected and post-fixed in 4%PFA at room temperature with rocking for 24h prior to paraffin embedding. Eight random mice per condition were chosen from the original cohort of mice for immunohistochemical analysis. Fixed tissue was process in a Donatello Automatic Sample Processor (Diapath) and embedded in paraffin. 2μm thick sections were collected on glass slides. For staining of the inguinal adipose tissue antigen retrieval with tri-sodium citrate (pH=6) was carried out for 30 min prior to a 1-h-incubation with the primary antibody at room temperature. The 2 steps HRP polymer Kit with DAB (DCX, PD000POL-K) was used for detection followed by hematoxylin counterstaining. Liver sections were stained with hematoxylin and eosin in an Erpedia Gemini AS strainer (Fisher Scientific).

Primary and secondary antibodies and concentrations are listed in Table S1. Processing and staining of adipose tissue and liver was performed by the Histology facility at the VBCF.

### Imaging

Images of brain immunofluorescence were acquired using an inverted Axio Observer microscope (ZEISS) equipped with a CSU X1 confocal scanning unit (ZEISS) and an EM-CCD camera (Hamamatsu, C9100-13). 40x and 63x oil Plan-Apochromat objective lenses (ZEISS) were used to image through whole 40 μm sections with a z-step of 1 μm. Images were stitched using ZEN Blue software (ZEISS).

Images of adipose tissue and liver were acquired using a Pannoramic FLASH 250 II (3DHISTECH) equipped with a CIS VVC FC60FR19CL camera (Vital Vision Technology) and a 20× Plan-Apochromat objective. Representative images were cropped using the CaseViewer software.

### Image quantifications

Quantifications of cells in the neurogenic lineage were performed using the Fiji software (Schindelin 2012) with the Cell Counter plugin. NSCs were identified by the localisation of a DAPI^+^ nucleus in the SGZ, the extension of a single GFAP^+^ vertical projection towards the molecular layer and the help of the YFP lineage tracing reporter (except in C57BL/6J mice), that is brighter in NSCs than other cells. IPCs were identified as Ki67^+^ cells in the SGZ. Neuroblasts were identified as DCX^+^ cells in the SGZ. Newly born neurons were identified as NeuN^+^ cells that colocalised with YFP (in GlastCreER^T2^;RYFP mice) or EdU (in C57BL/6J mice) in the SGZ and granule cell layer of the DG. In GlastCreER^T2^;RYFP mice, NeuN counts were normalised to the recombination rate. New cells in the DG were identified by EdU labelling. For cell numbers, 1 DG per sample was quantified (two for NSCs). The freehand line tool was used to measure the length of the SGZ, to which the total cell counts were normalised. For percentage of proliferating or EdU-labelled NSCs, at least 120 cells from two DG were counted in young mice, and 70 in older mice. All data were counted with group blinding within each experiment.

The appropriate sample size was based on previous publications studying NSC behaviour (Andersen 2014, Urban 2016, Blomfield 2019, Austin 2021), accounting for the increased variability of dietary interventions compared to genetic ones and considering female and male mice as separate groups. For the quantifications based on the YFP reporter, samples with a recombination rate lower than 80% were excluded. Samples with poor staining quality were also excluded from the quantifications.

### Statistical analysis

All statistical analyses were conducted using GraphPad Prism version 9.3.1 for macOS (GraphPad Software, San Diego, California, USA). Two-tailed unpaired student t tests were used for the comparison of two conditions. Whenever the samples did not pass a Shapiro-Wilk normality test (p<0.05), the non-parametric Mann-Whitney test was conducted (Figures S5A, B and S6A). A two-way ANOVA for repeated measures and multiple comparisons with a Šídák correction was performed to compare the weight of control and IF mice over time in Figure 1C. In Figure S2B, the weight gain data had to be fitted into a mixed effects model because of a missing value. For sex comparisons, a two-way ANOVA followed by Tuckey’s multiple comparisons test was performed. Error bars represent mean + standard deviation (SD). Significance is stated as follows: ns, p>0.05; *, p<0.05; **, p<0.01; ***, p<0.001; ****, p<0.0001. In graphs displaying weight (Figures 1C and S2B) and in Figures S1A, B and S7 p>0.05 is indicated by absence of significance symbol. Independent biological replicates are represented as dots in the bar plots.

## Acknowledgments

We thank all facilities at IMBA without which this work would have not been possible, in particular the animal house, bio-optics, histology and the workshop. We are particularly grateful to Mark Mattson, Seungjoon Park and Sandrine Thuret for sharing their experimental details with us. We want to specially thank Sandrine Thuret also for her helpful comments on our manuscript. We also thank all members of the Urbán Lab at IMBA for their critical reading of the manuscript.

## Author Contributions

R.G.S. co-developed the concept, performed experiments, analysed data, and co-wrote the manuscript. A.D., T.K. and I.C.E. performed experiments and edited the manuscript. N.U. conceived the original study, co-developed the concept, and co-wrote the manuscript.

---

**Figure S1.**
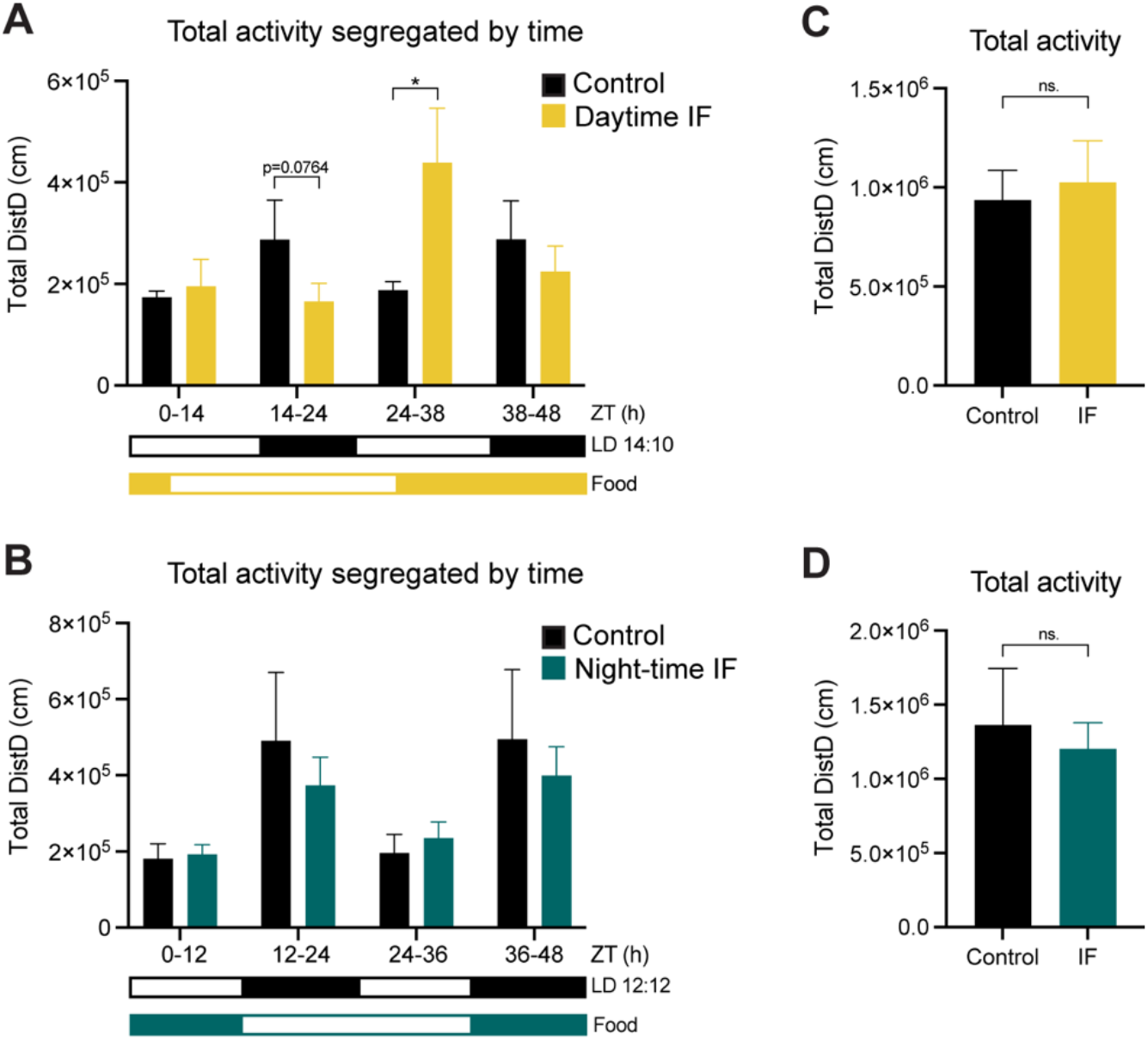
Total activity is not affected by day-nor night-time IF. (**A**, **B**) Total mouse locomotor activity displayed as sum of activity during 1 month of IF segregated by time of the day (grouped according to the light phase). Black and white boxes under the graphs indicate respectively the dark and light phases of the light:dark (LD) cycle with zeitgeber time (ZT) 0 being the time of lights ON. Yellow/blue and white boxes indicate the presence and absence of food respectively. Expected differences between light and dark phases were observed (not displayed in graph). Daytime IF increased locomotor activity during the feeding day while night-time IF preserved the activity pattern. (**C**,**D**) Total mouse locomotor activity during 1 month of diet shows no difference between IF an control mice regardless of the time of food administration. Black: control (ad libitum, n = 3 in daytime IF and n = 6 in night-time IF), yellow: daytime IF (n = 4), dark blue: night-time IF (n = 6). Bars and error bars represent mean + SD. Significance values: ns, p>0.05; ***, p<0.001.

**Figure S2.**
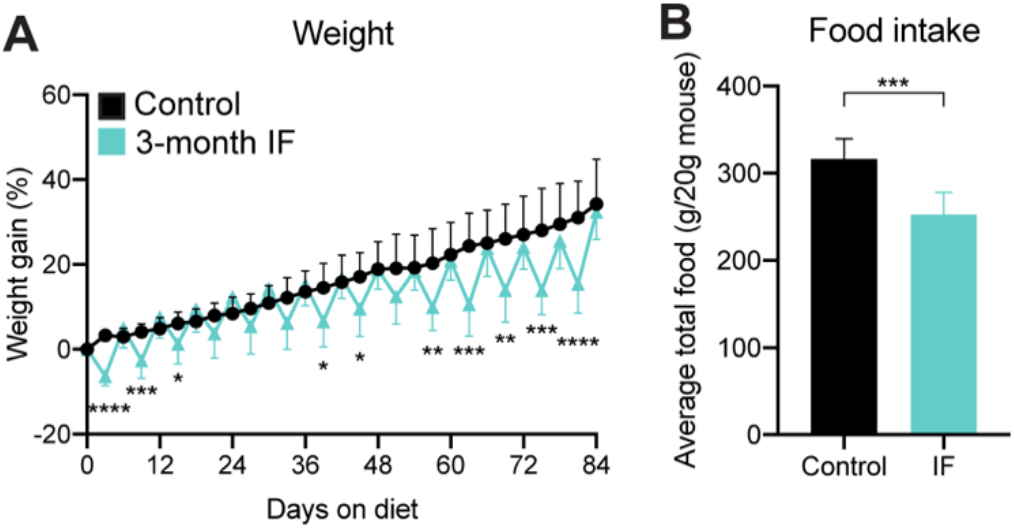
Mouse weight and food intake of mice during 3 months of IF. (**A**) Mouse weight shown as the percentage of weight difference to the first day of the diet. Weight oscillations persist throughout the 3 months of IF and there is no weight loss on refeeding days compared to control mice. n = 17. (**B**) Average total food consumed for 3 months shown as g of food per 20g of mouse weight (weight reference of mice at P65). n = 17. Significance values: no symbol, p>0.05; *, p<0.05; **, p<0.01; ***, p<0.001; ****, p<0.0001. Black: control (ad libitum), light blue: 3-month IF.

**Figure S3.**
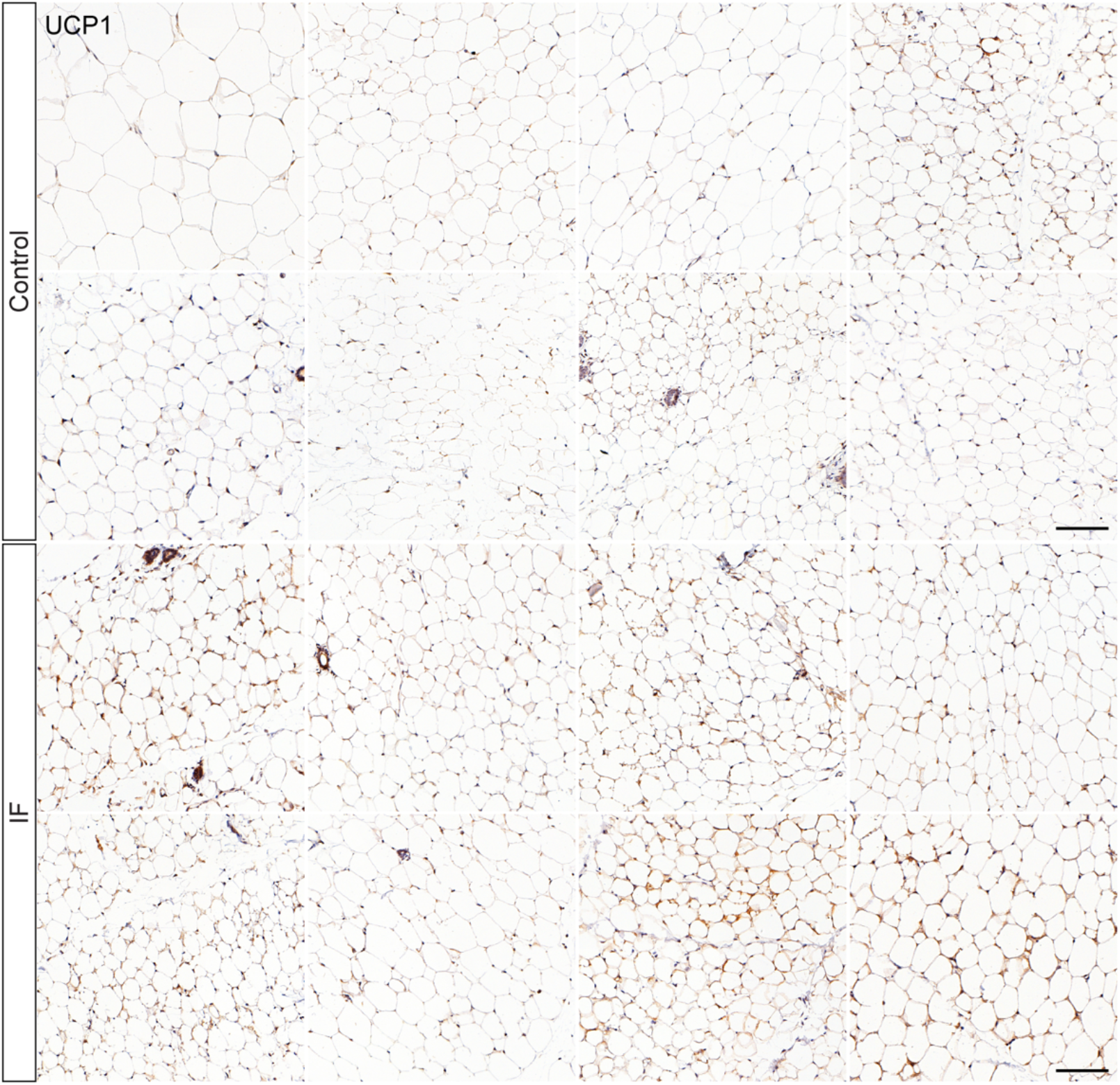
Night-time IF promotes mild and variable browning of inguinal adipose tissue. Images of inguinal adipose tissue stained for uncoupling protein 1 (UCP1) of control and mice that underwent 3 months of IF. 8 mice per condition were randomly chosen from the experiment shown in **Figure 3** for immunohistochemical evaluation. Each image corresponds to an individual mouse. Scale bar: 100μm.

**Figure S4.**
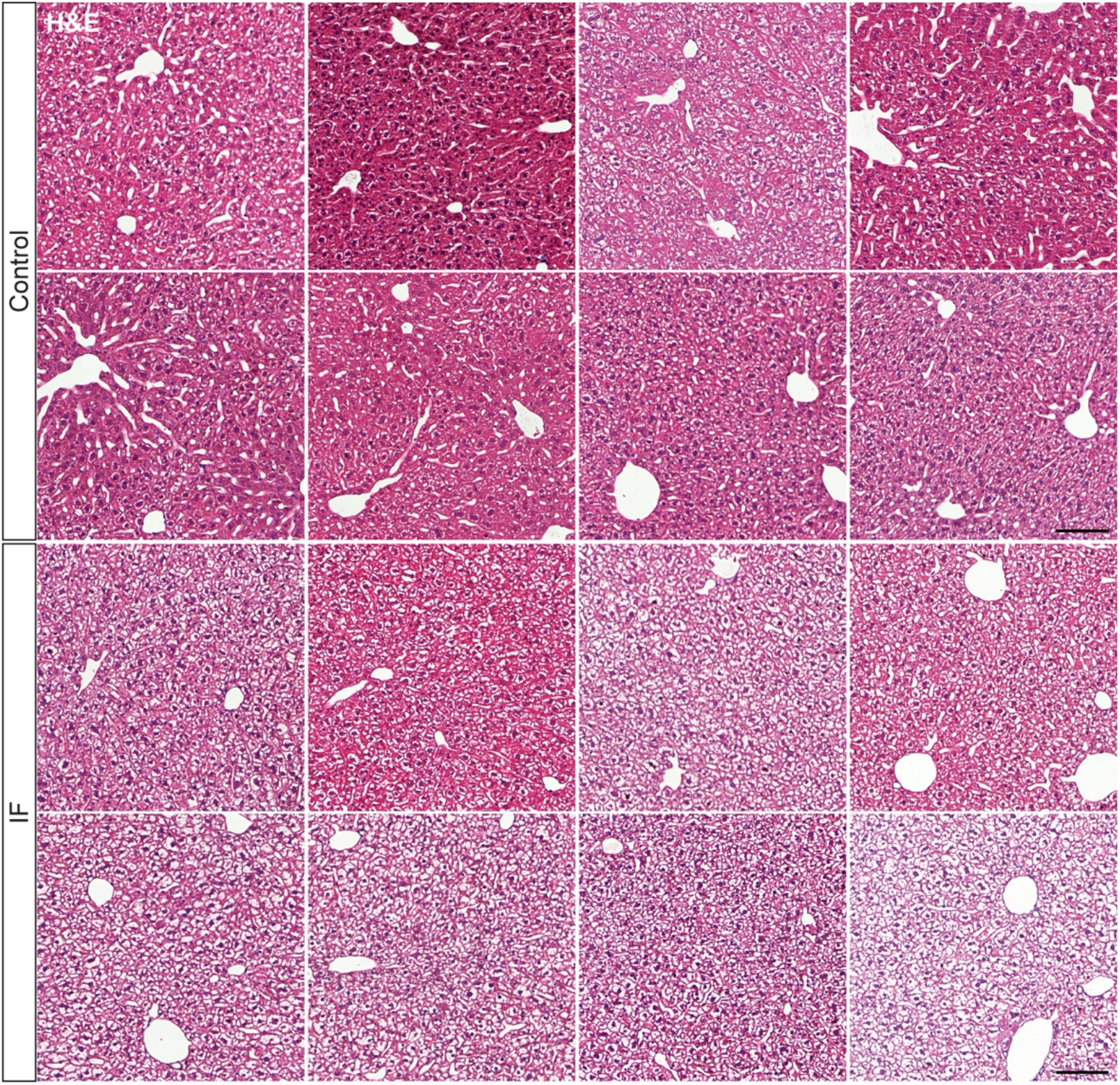
Hepatocytes display edematous morphology upon night-time IF. Images of H&E-stained liver of control and mice that underwent 3 months of IF. 8 mice per condition were randomly chosen from the experiment shown in **Figure 3** for immunohistochemical evaluation. Each image corresponds to an individual mouse. Scale bar: 100μm.

**Figure S5.**
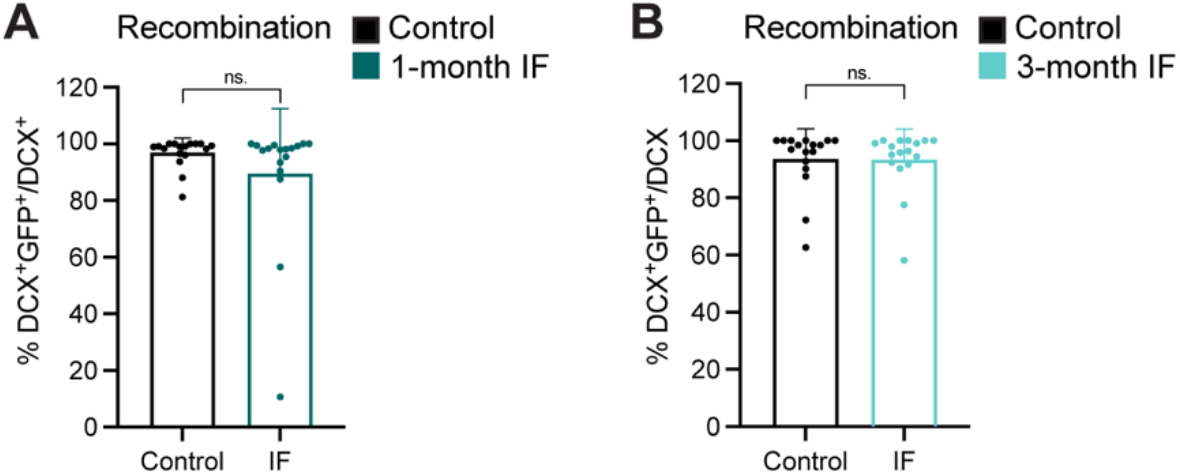
High levels of recombination in Glast-CreER^T2^;RYFP throughout experiments and treatments. (**A**, **B**) Recombination of the YFP reporter in Glast-CreER^T2^;RYFP upon tamoxifen induction shown as YFP^+^ neuroblasts over the total number of neuroblasts. (**A**) Recombination levels evaluated at the end of the 1-month treatment. (**B**) Recombination levels evaluated at the end of the 3-month treatment. Black: control (ad libitum) for each experiment, dark blue: 1-month IF, light blue: 3-month IF. Bars and error bars represent mean + SD; dots represent individual mice. Significance values: ns, p>0.05.

**Figure S6.**
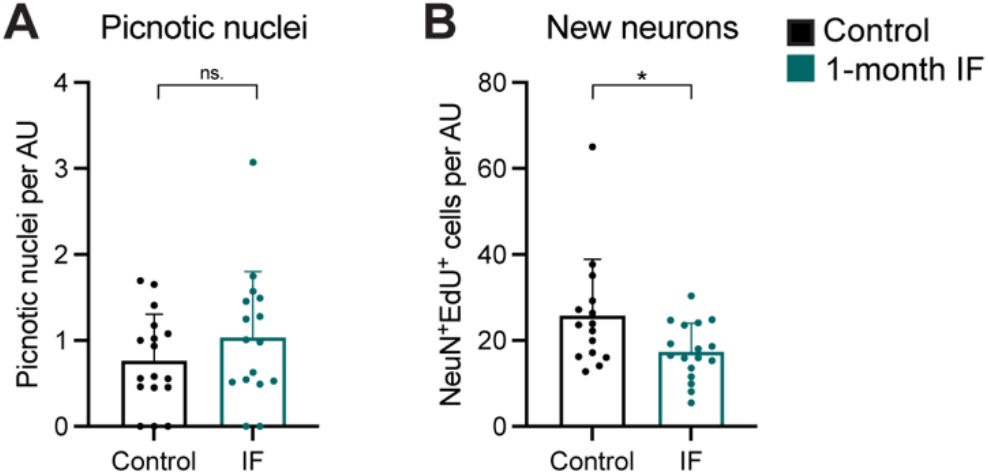
Transient decrease in the generation of new neurons upon 1 month of IF. See experimental design in **Figure 2A**. (**A**) Picnotic nuclei normalised to DG length as a proxy for cell death, which was not affected by IF. (**B**) Number of EdU-labelled neurons (as NeuN^+^ cells) after a 10-day chase, showing a decrease in the number of new neurons generated. Bars and error bars represent mean + SD; dots represent individual mice. Significance values: ns, p>0.05; *, p<0.05. Black: control (ad libitum), dark blue: IF.

**Figure S7.**
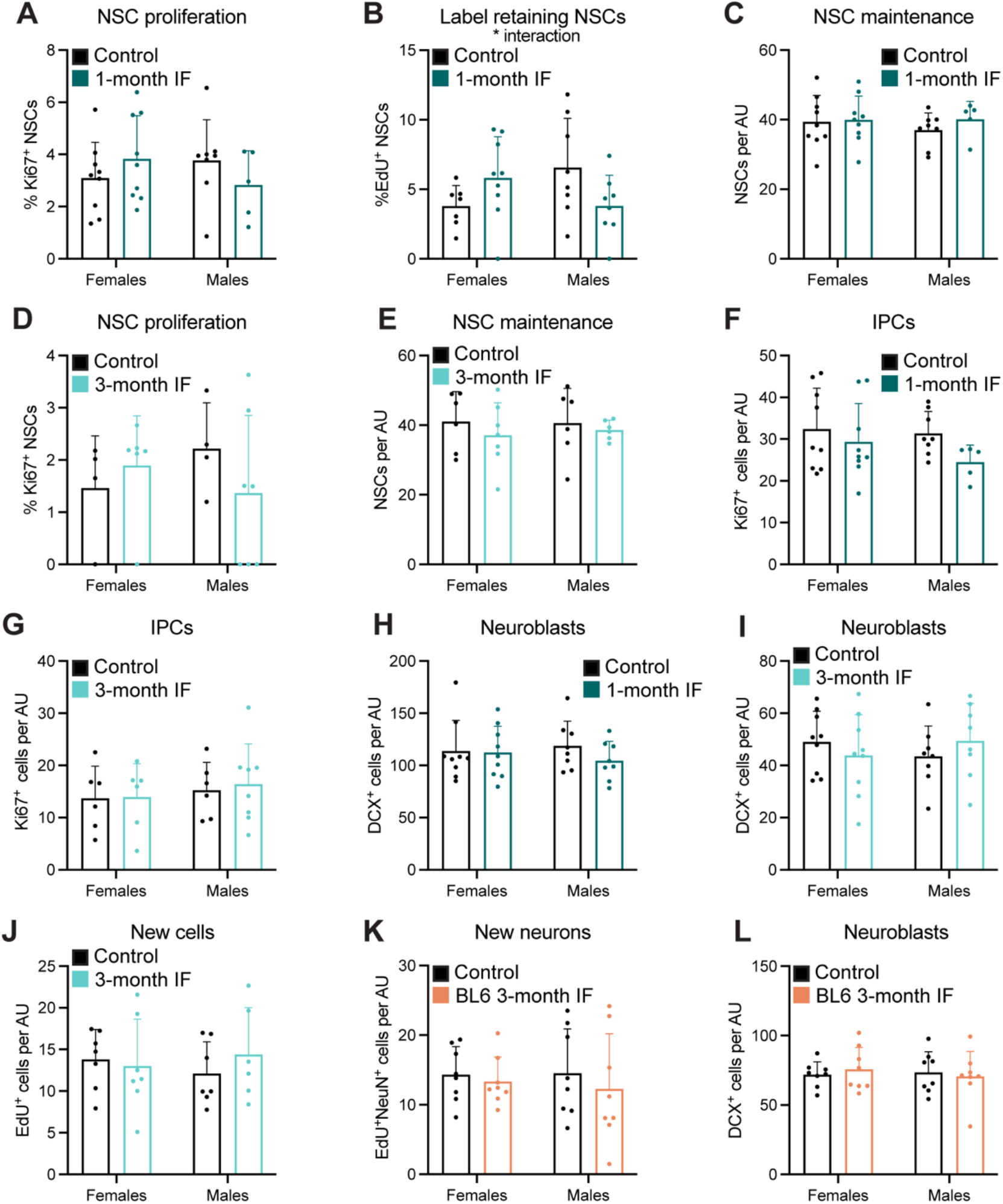
Male and female mice show similar responses to IF. (**A**-**L**) Data of all graphs in the main figures segregated by sex showing that the response of adult neurogenesis to IF is sex-independent. The percentage of label retaining NSCs after 1 month of IF (**B**) shows an interaction between diet and sex that is not translated in the following steps of the neurogenic lineage. Graphs are displayed in order of appearance on the text. Bars and error bars represent mean + SD; dots represent individual mice. Significance values: absence of sign, p>0.05; *, p<0.05. AU: arbitrary unit. Black: controls (ad libitum), dark blue: 1-month IF, light blue: 3-month IF, orange: C57BL/6J 3-month IF.

**Figure S8.**
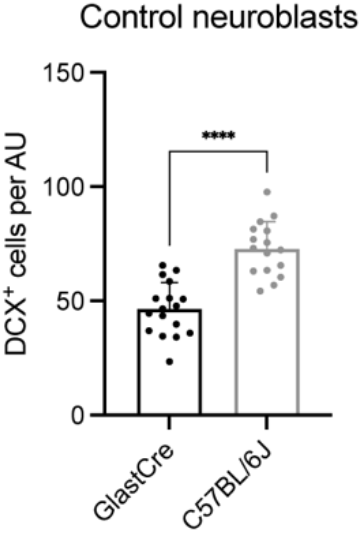
Differences in basal levels of neurogenesis between mouse strains. Neuroblasts (DCX^+^ cells) in Glast-CreER^T2^;RYFP and C57BL/6J mice. The neurogenic rate (shown as number of neuroblasts) is a 40% lower in 5-month-old Glast-CreER^T2^;RYFP mice with a mixed genetic background than in C57BL/6J mice of matched age. Bars and error bars represent mean + SD, dots represent individual mice. Significance values: *, p<0.0001. AU: arbitrary unit. Black: 5-month-old Glast-CreER^T2^;RYFP control, gray: 5-month-old C57BL/6J control.

**Table S1.** List of primary and secondary antibodies used for immunohistochemistry.

| **Target molecule** | **Species** | **Dilution** | **Company** | **Catalog number** |
| --- | --- | --- | --- | --- |
| GFP | chicken | 1:5000 | Abcam | ab13970 |
| GFAP | rabbit | 1:500 | DAKO | Z033401-2 |
| GFAP | rat | 1:500 | Invitrogen | 130300 |
| Ki67 | rat | 1:200 | eBioscience | 14-5698-82 |
| DCX | rabbit | 1:1000 | Abcam | ab18723 |
| NeuN | mouse | 1:1000 | Chemicon | MAB377 |
| UCP1 | rabbit | 1:1000 | Abcam | ab10983 |
| Chicken IgG | donkey – 488 | 1:500 | Jackson Laboratories | 703-545-155 |
| Rabbit IgG | donkey – Cy3 | 1:500 | Jackson Laboratories | 711-166-152 |
| Rat IgG | donkey – Cy3 | 1:500 | Jackson Laboratories | 712-166-153 |
| Rat IgG | donkey – 647 | 1:500 | Jackson Laboratories | 712-605-150 |
| Mouse IgG | donkey – 647 | 1:500 | Jackson Laboratories | 715-606-151 |

